# Population genomics of wall lizards reflects the dynamic history of the Mediterranean Basin

**DOI:** 10.1101/2021.05.26.445763

**Authors:** Weizhao Yang, Nathalie Feiner, Daniele Salvi, Hanna Laakkonen, Daniel Jablonski, Catarina Pinho, Miguel A. Carretero, Roberto Sacchi, Marco A. L. Zuffi, Stefano Scali, Konstantinos Plavos, Panayiotis Pafilis, Nikos Poulakakis, Petros Lymberakis, David Jandzik, Ulrich Schulte, Fabien Aubret, Arnaud Badiane, Guillem Perez i de Lanuza, Javier Abalos, Geoffrey M. While, Tobias Uller

## Abstract

The Mediterranean Basin has experienced extensive change in geology and climate over the past six million years. Yet, the relative importance of key geological events for the distribution and genetic structure of the Mediterranean fauna remains poorly understood. Here, we use population genomic and phylogenomic analyses to establish the evolutionary history and genetic structure of common wall lizards (*Podarcis muralis*). This species is particularly informative because, in contrast to other Mediterranean lizards, it is widespread across the Iberian, Italian, and Balkan peninsulas, and in extra-Mediterranean regions. We found strong support for six major lineages within *P. muralis*, which were largely discordant with the phylogenetic relationship of mitochondrial DNA. The most recent common ancestor of extant *P. muralis* was likely distributed in the Italian Peninsula, and experienced an “Out-of-Italy” expansion following the Messinian salinity crisis (~5 Mya), resulting in the differentiation into the extant lineages on the Iberian, Italian and Balkan peninsulas. Introgression analysis revealed that both inter- and intraspecific gene flow have been pervasive throughout the evolutionary history of *P. muralis*. For example, the Southern Italy lineage has a hybrid origin, formed through admixture between the Central Italy lineage and an ancient lineage that was the sister to all other *P. muralis*. More recent genetic differentiation is associated with the onset of the Quaternary glaciations, which influenced population dynamics and genetic diversity of contemporary lineages. These results demonstrate the pervasive role of Mediterranean geology and climate for the evolutionary history and population genetic structure of extant species.

## Introduction

The reconstruction of evolutionary history is essential if we are to understand the factors and processes that explain contemporary patterns of biodiversity (Avise 2000). The Mediterranean Basin is considered a biodiversity hotspot because of its species richness and high degree of endemism (Médail and Quezel 1997; Myers et al. 2000). However, the processes responsible for Mediterranean diversity and biogeography have proven difficult to resolve because of the complex geological and climatic history of this region (Cavazza and Wezel 2003; Médail and Diadema 2009; Thompson 2005; Hewitt, 2011a). Defining events include the Messinian salinity crisis (MSC; 5.96-5.33 million years ago, Mya; Krijgsman et al. 1999, 2010; Duggen et al. 2003), a period of progressive aridity that led to the extinction of subtropical Tertiary lineages and the diversification of arid-adapted lineages as well as extensive faunal exchange within the Mediterranean and with Africa (Jiménez-Moreno et al. 2010; Fiz-Palacios and Valcárcel 2013). The refilling of the Mediterranean (García-Castellanos et al. 2009) resulted in the isolation of biota on islands and peninsulas, and the onset of the Mediterranean climate of today (3.4-2.8 Mya) significantly changed ecological communities (Tzedakis 2007; Postigo et al. 2009). The later Quaternary climatic oscillations (starting ca. 2.5 Mya), characterized by the alternation of colder (glacial) and warmer (interglacial) periods, further affected the distribution of many species due to recurrent range shifts with associated cycles of demographic expansion-contraction (Hewitt 1996, 2000; Provan and Bennet 2008; Taberlet et al. 1998). During glacial periods, the Mediterranean Basin provided refugial areas, where the long-term persistence of isolated populations frequently resulted in the formation of new allopatric lineages (Hewitt 1996, 2004; Wiens 2004; Gentili et al. 2015; Mairal et al. 2017).

While all these events contributed to the evolution of the Mediterranean fauna, their respective roles for explaining the genetic structure and geographic distribution of extant taxa remain poorly understood (Hewitt 2011a). One reason for this is that reconstruction of evolutionary history can be challenging, especially when lineages have been subject to repeated range expansions and contractions over the Quaternary climatic cycles (e.g., Buckley 2009; Hewitt 2011b; Carstens et al. 2013). Introgression imposes further challenges for reconstructing historical relationships among taxa, as well as estimates of taxonomic diversity (e.g., Naciri and Linder 2015; Mallet et al. 2016). Approaches based on representative sampling of genome-wide genetic variability allow powerful and refined phylogeographical inference overcoming many of the limitations of more traditional molecular markers (Delsuc et al. 2005; Cutter 2013; McCormack et al. 2013). Such phylogenomic approaches can be particularly useful in resolving evolutionary affinities in situations where clades are separated with short internal branches (Pollard et al. 2006) or where hybridization has occurred (Cui et al. 2013; McCluskey and Postlethwait 2015; Malinsky et al. 2018; Chen et al. 2019).

Wall lizards of the genus *Podarcis* are currently represented by 24 to 25 species (Speybroeck et al. 2020) and are a characteristic fauna of the Mediterranean Basin and its islands. Similar to many other Mediterranean animals (reviewed in Hewitt 2011a), wall lizards commonly exhibit high regional endemism, with species typically being restricted to one of the Balkan, Iberian, and Italian peninsulas, or one or several Mediterranean islands (Poulakakis et al. 2005; Psonis et al. 2021; Salvi et al. 2021; Yang et al. 2021). A notable exception is the common wall lizard (*P. muralis*). This species is not only widespread, being distributed from Iberia to Asia Minor, but it is also native to extra-Mediterranean regions in Western, Central and Eastern Europe (Schulte et al. 2008; fig. 1A). Previous studies based on DNA sequence data suggest that such widespread geographic distribution has been accompanied by regional differentiation into more than 20 genetic lineages that purportedly diverged during the Pleistocene glaciations (Salvi et al. 2013). These lineages were defined by divergence in mitochondrial DNA, several of them separated by low genetic divergence (i.e. short internal branches), and their phylogenetic relationships therefore remain largely unresolved (Salvi et al., 2013). Moreover, recent analyses of single nucleotide variants (SNV) data have demonstrated extensive gene flow even between distantly related mtDNA lineages of *P. muralis* (Yang et al. 2018; 2020). This suggests a much more complex scenario than what can be revealed by traditional phylogenetic studies.

**Fig. 1.**
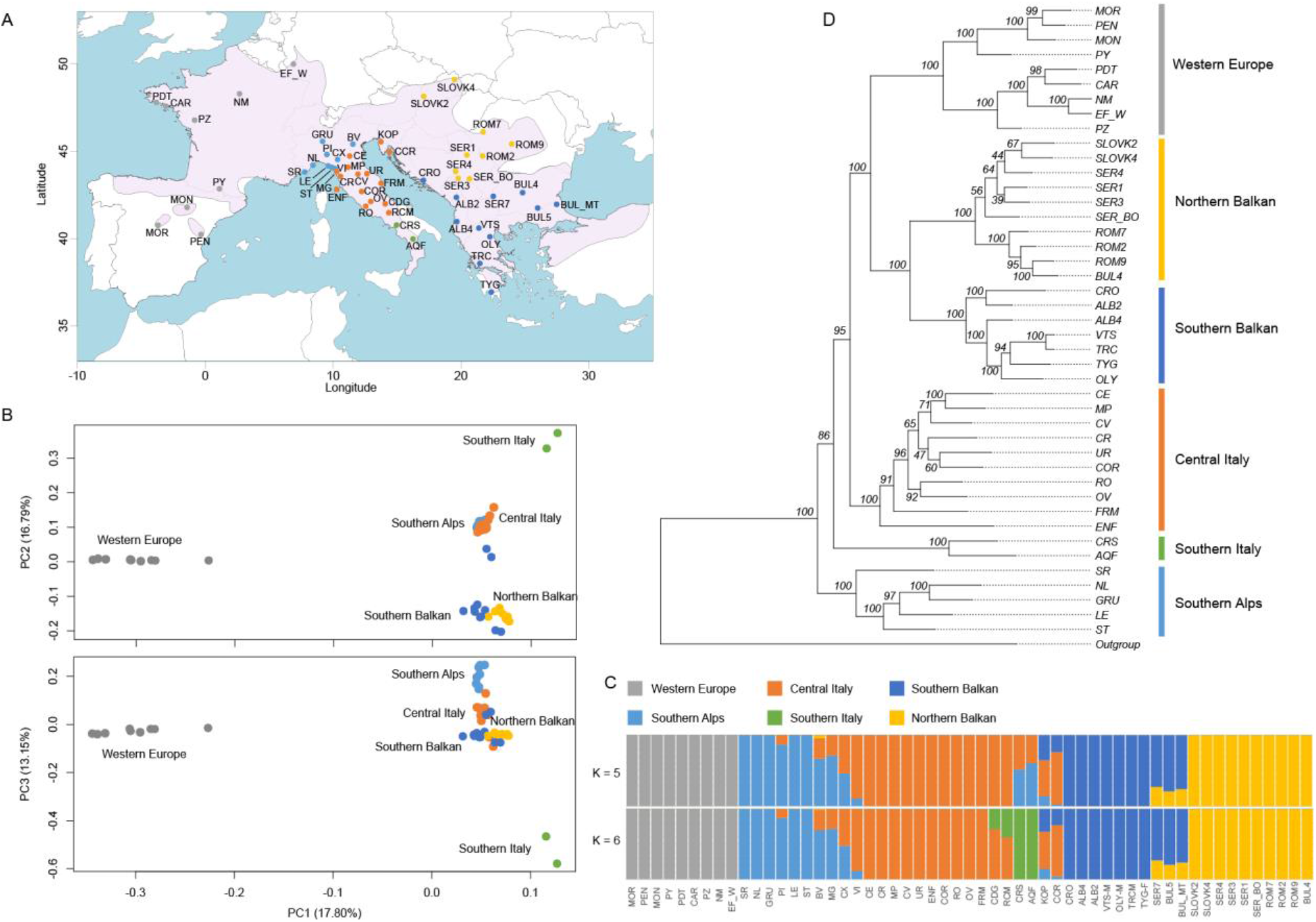
Population genetic and phylogenetic analyses of *P. muralis*. (A) The sampling localities for the RAD-Seq part of this study. The pink area represents the distribution range of *P. muralis.* For sampling locations of WGS samples, see supplementary fig. S1. (B) PCA plots of genetic distance for all 55 individuals based on RAD-Seq data. (C) Admixture clustering of individuals into five and six groups (K). The proportion of each individual’s genome assigned to each cluster is shown by the length of the colored segments. (D) Maximum likelihood phylogeny inferred based on RAD-Seq data. The numbers above branches indicate the bootstrap values for each node. For all panels, genetic lineages are color-coded consistently.

In this study, we implemented a phylogenomic approach based on both restriction site-associated DNA sequencing (RAD-Seq) and whole genome sequencing (WGS) to identify the processes that have shaped genetic differentiation and biogeography of *P. muralis.* We had three specific aims. First, to identify lineages of *P. muralis* from both nuclear and mitochondrial genomic data and establish their phylogenetic relationships. Second, to establish evidence for hybridization and introgression between each of the major *P. muralis* lineages and between *P. muralis* and other *Podarcis* species that currently overlap in their distribution. Third, to reconstruct the biogeographic and demographic history of *P. muralis* in Europe, and assess whether these reflect the paleogeographic and climatic history of the Mediterranean Basin.

## Results

We collected samples from 55 locations for RAD-Seq covering the previously suggested genetic lineages within *P. muralis* (fig. 1A; supplementary table S1; Gassert et al. 2013; Salvi et al. 2013; Jablonski et al. 2019). In total, 28,039 SNVs were obtained with a mean coverage of 17.10 per site and an average genotyping rate of 0.95. In addition, whole genomes of 16 individuals were sequenced (supplementary fig. S1; supplementary table S2), obtaining 9,699,080 nuclear SNVs with a mean coverage of 11 and genotyping rate above 0.99. We also sequenced the full mitochondrial genomes of those individuals (13,884 bp). One individual of *P. bocagei*, *P. siculus*, and *P. tiliguerta* served as outgroups for all analyses, and *Archaeolacerta bedriagae* was further included as outgroup for WGS data (see Yang et al. 2021 for a phylogenomic analysis of all *Podarcis* species). In addition, we also included WGS data of 13 additional *Podarcis* lineages (Yang et al. 2021; supplementary table S3) to assess gene flow between *P. muralis* and other *Podarcis* species.

### Population structure analysis

We inferred the genetic relationship between the *P. muralis* individuals based on genotypes from RAD-Seq data using a principal component analysis (PCA; Chang et al. 2015) and ADMIXTURE clustering (Alexander et al. 2009). Results of both PCA and ADMIXTURE supported distinct *P. muralis* lineages with a strong geographic structure. In the PCA, the first, second and third principal components (variances explained: 17.80%, 16.79% and 13.15%, resp.) largely separated populations into five different lineages: Western Europe (WE), Balkan, Southern Italy (SI), Southern Alps (SA) and Central Italy (CI) (fig. 1B). In the ADMIXTURE clustering, the best supported number of presumed ancestral populations (K = 5) further divided the Balkan populations into Southern Balkan (SB) and Northern Balkan (NB). The lineage in Southern Italy, SI, showed an admixed pattern of the Central and Northern Italian lineages, CI and SA, at K = 5, but was recovered as an independent cluster (SI) at K = 6 (fig. 1C). Several other admixed populations were also identified by this population structure analysis, including admixture between three lineages, CI, SA, and SB, on the Balkan peninsula (locations CCR and KOP), and between SA and CI in Northern Italy (locations BV, CE, and MG) (fig. 1C).

### Phylogenetic relationship between lineages

Following the population genetic structure, we inferred the phylogenetic relationship between the six major lineages (CI, NB, SA, SB, SI, and WE). Concatenated maximum likelihood (ML) analysis yielded a highly resolved phylogeny based on RAD-Seq data: 80% of the branches exhibited bootstrap values > 80%, and the branches leading to the six major lineages showed bootstrap values equal to 100% (fig. 1D). In this phylogeny, SA formed the sister clade to all other lineages, CI formed the sister clade to the WE and the Balkan clade, where each of the latter two were divided into northern and southern sub-clades (note that the subdivision of WE into northern and southern sub-clades was not detected by the ADMIXTURE clustering). The only poorly supported node connecting the major lineages (bootstrap of 85%) was associated with the phylogenetic position of SI as a sister clade to all non-SA lineages. Considering this uncertainty and the admixed pattern of SI in the ADMIXTURE analysis, we also reconstructed the phylogeny by excluding all SI individuals, in which the phylogenetic relationship between all other lineages remained the same (supplementary fig. S2).

Next, we inferred the ML phylogeny based on WGS data for 16 individuals representing all major distribution ranges (Supplementary fig. S1). This WGS phylogeny was supported by bootstrap values of 100% for all nodes and its topology showed the same relationships among the six major lineages as the RAD-Seq phylogeny (Supplementary fig. S3). One individual from the admixed population KOP (fig. 1C), clustered with the Balkan lineages and was excluded from the following analyses (fig. 2). We also validated the phylogeny using a multispecies coalescent approach in ASTRAL-III (Zhang et al. 2018) based on local ML trees of 200 kb windows across the genome, and again obtained the same topology with bootstrap values of 100% for all branches (Supplementary fig. S4). These results congruently indicated that the obtained phylogeny was highly robust.

**Fig. 2.**
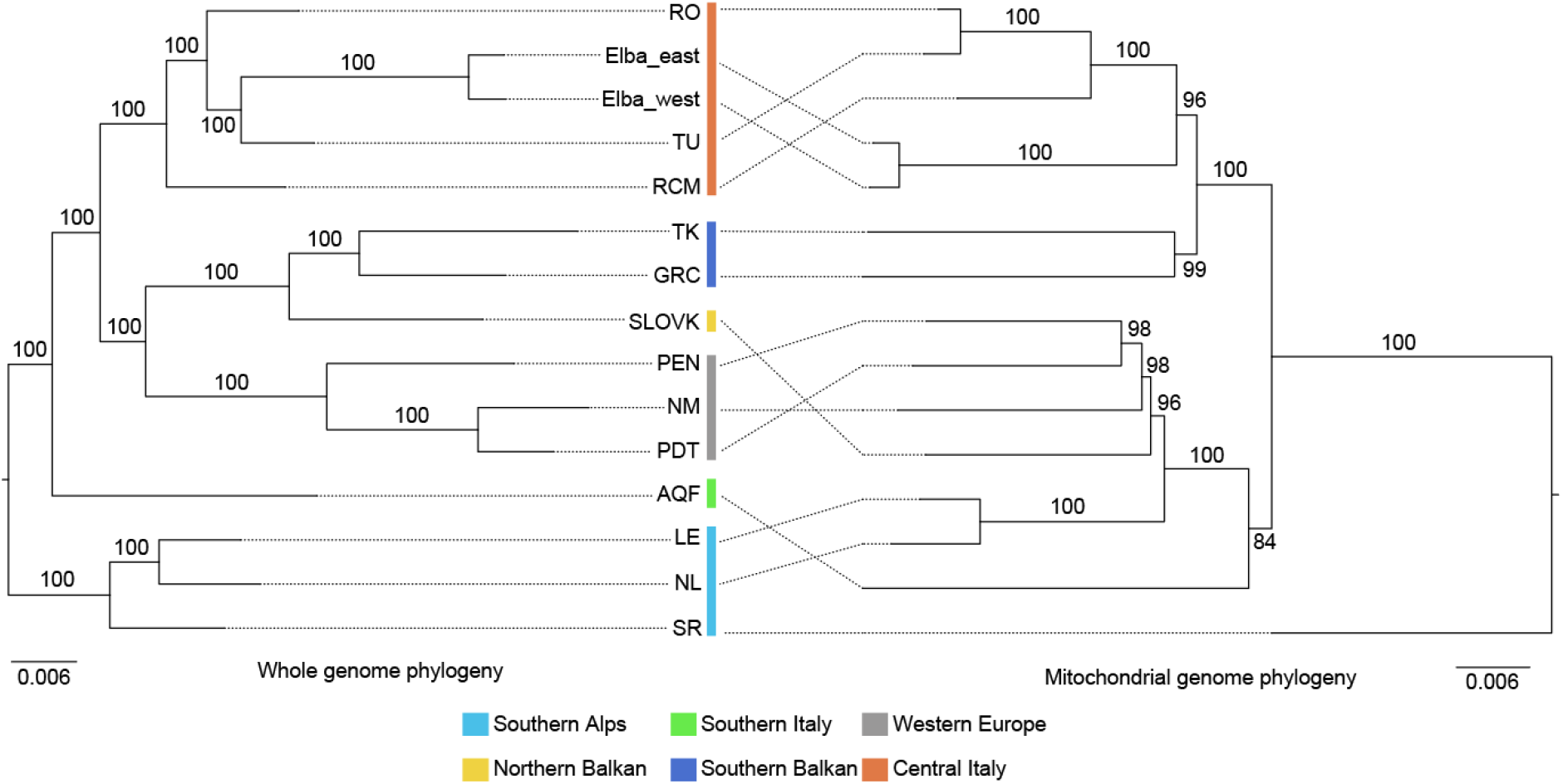
Mito-nuclear discordance for *P. muralis* lineages. The discrepancies between phylogenies based on whole-genome sequencing data (left) and mitochondrial genome data (right). Bootstrap values are provided for all nodes. Colors refer to genetic lineages.

We obtained an ML tree based on mitochondrial genome sequences with well supported clades (average bootstrap value > 97.57%; fig. 2). The mtDNA phylogeny was extensively discordant with the phylogeny derived from nuclear data (nuDNA). For example, the SA lineage was grouped with WE and SB in the mtDNA tree (which in turn grouped with the SI mtDNA lineage), while the other major mtDNA clade was formed by only CI and NB. Surprisingly, the mitochondrial genome of the individual from San Remo (SR) was not nested within the SA lineage, but was basal to all other lineages. Other mtDNA-nuDNA discordances were found for populations from Elba and in the contact zone between the SA and SB lineages on the Balkan Peninsula (fig. 2). We confirmed that the mitochondrial genomes of all the *P. muralis* lineages clustered as a monophyletic clade by including mitochondrial genomes of all the13 *Podarcis* species in the phylogenetic analysis (Supplementary fig. S5).

### Gene flow analysis

We first tested for interspecific gene flow between *P. muralis* lineages and the 13 *Podarcis* species and lineages (see Yang et al. 2021), using D statistics (supplementary table S3; Patterson et al. 2012). A total of 273 tests were performed, in which 127 (46.52%) significantly deviated from neutrality (Z-score > 3.3). The top 50, and more than 2/3 of all significant tests, involved the WE lineage, which excessively shared alleles with the common ancestor of the Iberian species group of *Podarcis* (here represented by *P. bocagei* and *P. hispanicus*) and the Ibiza wall lizard (*P. pityusensis*). Other signatures of interspecific gene flow were found between *P. siculus* and the CI and SA lineages, but the D and Z-scores were substantially lower than for WE and the Iberian *Podarcis* species (fig. 3A).

**Fig. 3.**
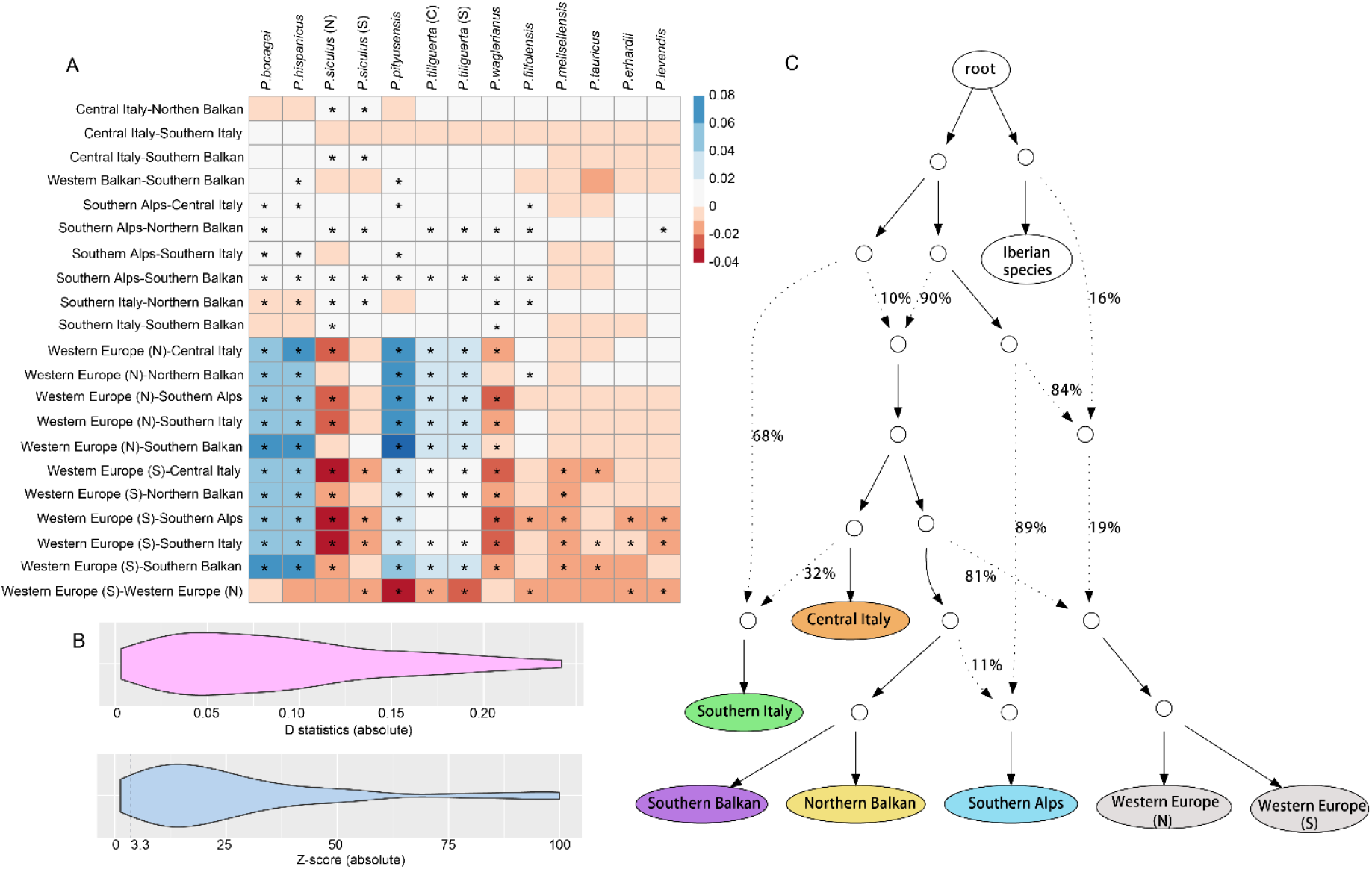
Analyses of gene flow for *P. muralis* lineages. (A) Interspecific introgression analyses between *P. muralis* and other *Podarcis* species using four-taxon D statistics with the test (muralis_A, muralis_B, target_taxon, outgroup). The lineage pairs of *P. muralis* are listed on the Y axis, and the targeted non-*muralis* species are listed on the X axis. The coloration of each square represents the D statistics, and the asterisk (*) indicates significant deviations from neutrality based on z-scores. (B) Distribution of D statistics and z-scores of intraspecific introgression analysis between *P. muralis* lineages. The result showed that most tests significantly deviated from neutrality. We list detailed information in supplementary table S4. (C) Admixture graph of *P. muralis* generated by qpGraph. Solid lines with arrows indicate tree-like evolution, whereas dash lines with arrows indicate admixture events. The numbers next to branches represent the proportion of alleles from a parental nodes.

Next, we investigated introgression within *P. muralis*. The results of D statistics revealed that 33 out of 35 tests (94.29%) significantly deviated from neutrality (Z-score > 3.3), suggesting substantial genetic exchange between the major *P. muralis* lineages (fig. 3B; detailed information in supplementary table S4). This pattern was supported by further analyses using phyloNet (supplementary fig. S6; Wen et al. 2008) and qpGraph (fig. 3C; Patterson et al. 2012). The phylogenetic network indicated a complex evolutionary history for the WE lineage, and demonstrated that the introgression from Iberian species (i.e., *P. hispanicus* complex) likely happened well before the WE lineage split into a northern and southern clade. This analysis also revealed that the SA lineage experienced introgression from the common ancestor of the two Balkan lineages. A complex history was inferred for the SI lineage, which received one part of its genome (phyloNet: 40%; qpGraph: 32%) from the CI lineage, and the other part (phyloNet: 60%; qpGraph: 68%) from a sister clade to all other *P. muralis* lineages (fig. 3C, supplementary fig. S6). This early diverging and extinct clade also contributed alleles (phyloNet: 44%; qpGraph: 10%) to the common ancestor of all extant lineages, except SA.

### The evolutionary history of the Southern Italy lineage

Gene flow analysis suggested that the current SI lineage resulted from the fusion of populations belonging to the CI lineage and an ancient SI lineage that was sister to all other lineages. Thus, the placement of the SI lineage in our inferred phylogeny was not supported by the phylogenetic network with gene flow (compare fig. 1D, fig. 2 and fig. 3C). To clarify the evolutionary history of the SI lineage, we estimated the distribution of different tree topologies across the genome based on 200 kb windows. A total of 5,576 high-quality local trees were inferred with average bootstrap value > 60%. Among these trees, 1,816 trees (32.57%) supported a monophyletic clade formed by the CI and SI lineages (Topology 3, 4 and 6 in fig. 4A&B). The windows supporting this relationship are referred to as the “CI-ancestry genome” of the SI lineage.

**Fig. 4.**
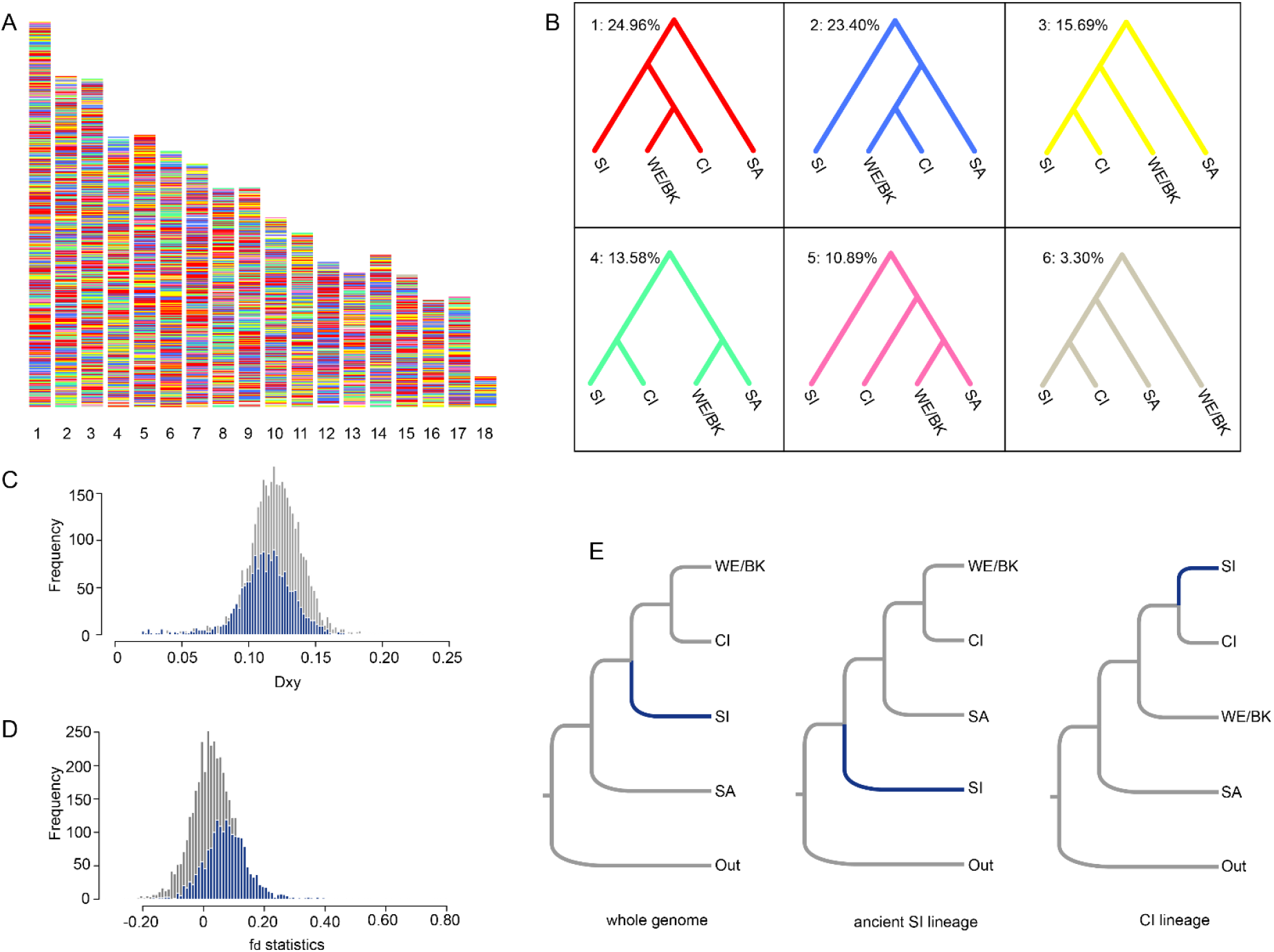
The ancestry and phylogenetic position of the Southern Italy lineage. (A) The distribution of local tree topologies across the genome. (B) The six most common topologies with the corresponding coloration as in panel (A). The value on the top left is the percentage of all 200 kb windows supporting the specific topology. (C) The distribution of Dxy between the two lineages for the CI-ancestry part of the genome (blue) and the rest of the genome (grey). (D) The distribution of fd statistics between the two lineages for the CI-ancestry genome (blue) and the rest of the genome (grey). (E) Multispecies coalescent tree topologies for distinct portions of the genome: whole genome, the ancient SI lineage part and the CI-ancestry part of the genome.

To test if introgression contributed to the discordance of lineage phylogeny and local trees besides incomplete lineage sorting, we further calculated the absolute genetic distance (Dxy) between CI and SI, and the fd-statistics (Martin et al. 2015) on the topology (WE/Balkan, CI, SI, Outgroup) for the CI-ancestry part and the rest of the genome. The Dxy of the CI-ancestry genome (0.1127) was significantly lower than the Dxy of the rest of genome (0.1198; p-value < 0.001; 1,000 iterations of permutation test; fig. 4C). Conversely, the fd of the CI-ancestry genome (0.0778) was significantly greater than that of the rest of genome (0.0265; p-value < 0.001; 1,000 iterations of permutation test; fig. 4D). These results strongly suggest that the CI-ancestry part of the SI genome was derived from an introgression between the CI and an ancient SI lineage.

Following these results, we next inferred the phylogeny using the same multispecies coalescent method but keeping the CI and the ancient SI lineage parts of the genome separated. The CI-ancestry genome of the SI lineage indeed formed a sister taxon with CI close to Western Europe / Balkan clade in the phylogeny of *P. muralis*. However, the ancient SI lineage formed an independent clade that branched off before any of the other *P. muralis* lineages (fig. 4E). This was consistent with the topology inferred by introgression analysis (fig. 3C), which indicated that about 68% of alleles in the genome of the SI lineage introgressed from this early diverged ancient SI lineage, and 32% of alleles introgressed from the CI lineage.

### Divergence time estimation

Divergence time estimation between *P. muralis* lineages was performed based on the 200-kb genomic windows (N = 346) for which local trees supported the topology derived from the ancient SI part of the genome (see fig. 4E). Two secondary calibrations were used from a fossil-calibrated Lacertini phylogeny of Garcia-Porta et al. (2019) - the root node (37.55 Mya) and the crown node of *Podarcis* (18.60 Mya). The time-calibrated tree (fig. 5A) revealed an early split of the ancient SI lineage estimated at ca. 6.24 Mya, followed by the separation of the SA lineage and the MRCA of all other lineages during the Messinian salinity crisis at ca. 5.76 Mya. The CI lineage separated shortly afterwards (ca. 4.90 Mya). The divergence between WE and the Balkan lineages was estimated to be ca. 4.05 Mya, and the divergences within the Western Europe and Balkan lineages were almost coinciding (ca. 2.54 Mya and ca. 2.58 Mya). We also estimated the divergence times based on the phylogeny without the ancient SI lineage, using the same methods, which generated consistent results (supplementary fig. S7).

**Fig. 5.**
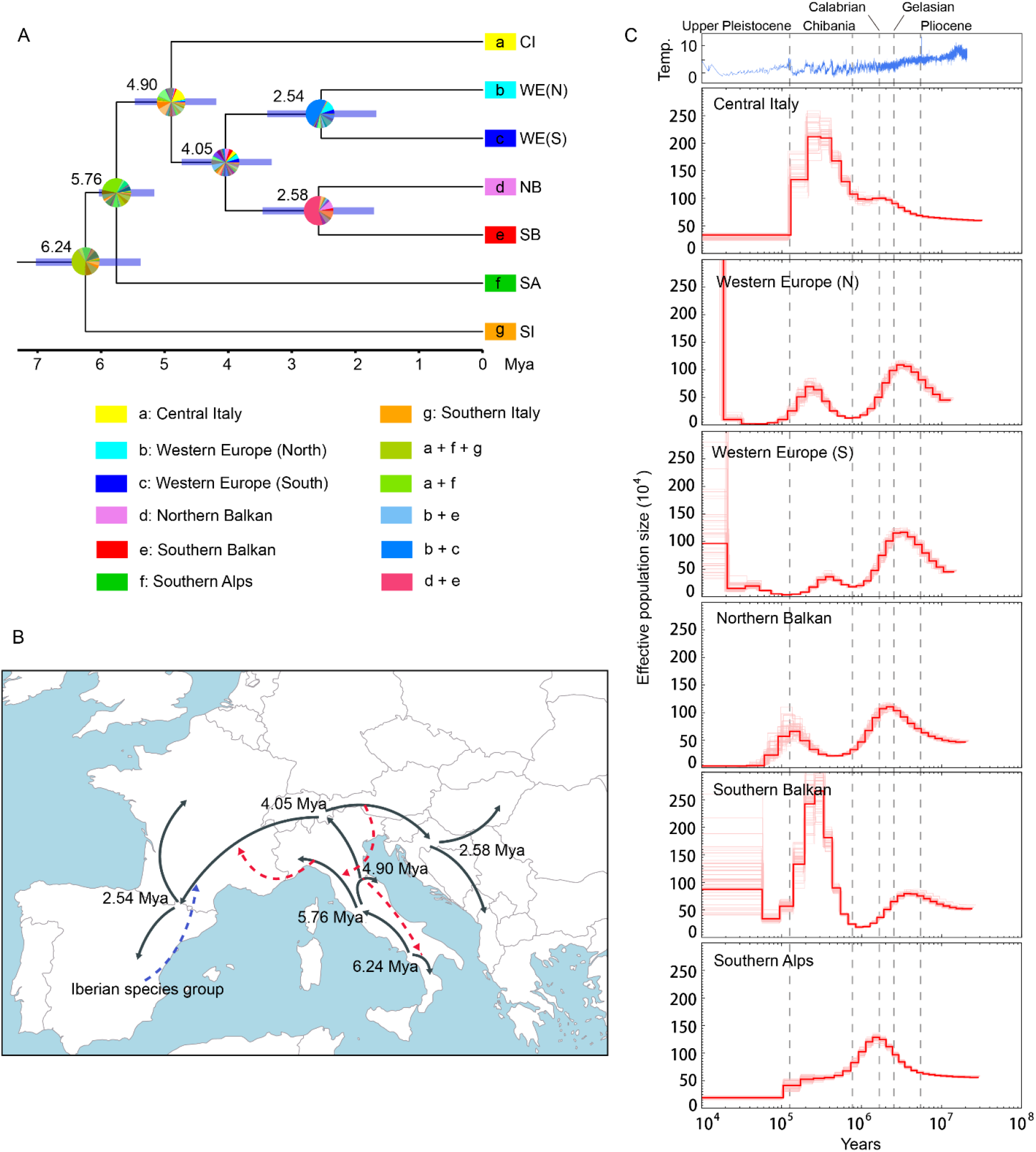
Biogeography and demographic dynamics for *P. muralis*. (A) Time-calibrated phylogeny for *P. muralis* lineages with ancestral area reconstructions. The numbers indicate the estimated ages for each node. The blue bars represent the confidence intervals of divergence times. The colored squares at tip nodes represent the current distribution ranges of lineages in seven biogeographic regions, and the pie charts indicate the proportional posterior probability of ancestral ranges inferred by the best-fitting model DIVALIKE. (B) Illustration of the inferred biogeographic history of *P. muralis* on a map of contemporary Europe. The solid lines with arrows indicate the tree-like divergence and dispersal. Dashed lines with arrows indicate interspecific introgression (blue) or intraspecific introgression (red). (C) Demographic history of five major *P. muralis* lineages (note that both Northern and Southern WE are shown in separate panels). Effective population size over time was estimated using the pairwise sequential Markovian coalescent model. The red lines represent the population dynamics, and the light red lines represent the results of 100 bootstrap replicates. The dash lines indicate the start of geological periods. The top panel shows the dynamics of global temperature.

### Estimation of biogeographic and demographic history

To infer the possible ancestral distribution range and the biogeographical events leading to the extant distribution of *P. muralis* lineages, we used BioGeoBEARS (Matzke 2013) with a total of three models - dispersal extinction cladogenesis (DEC), dispersal vicariance analysis (DIVALIKE), Bayesian inference of historical biogeography for discrete areas (BAYAREALIKE). We defined seven biogeographic areas: Iberian Peninsula, Western Europe, Southern Alps, Central Italy, Southern Italy, Northern Balkan and Southern Balkan. The results showed that the DIVALIKE was the best-fitting model (AICc = 50.69; supplementary table S5). All three models suggested very similar patterns, in which the ancestral range of *P. muralis* was located on the Italian Peninsula. The three major lineages found there today (the SI, SA and CI lineages) appear to have separated within the Italian Peninsula, followed by an out-of-Italy dispersal towards the Balkan and Iberian Peninsulas, and into north-western and central Europe (fig. 5A,B; supplementary fig. S8).

We further reconstructed the detailed demographic history of each lineage using the pairwise sequential Markovian coalescence (PSMC; Li and Durbin 2011; see fig. 5). The SI lineage was excluded in this analysis due to its highly heterogeneous genome. With respect to the demographic history, the PSMC model suggested an initial population expansion for all lineages until the Lower Pleistocene (ca. 2 Mya; fig 5). Subsequently, all lineages except CI experienced a population decline at the beginning of the Quaternary climate oscillations (Gelassian and Calabrian). The CI, NB, SB and WE lineages then experienced another population expansion from ca. 700 thousand years ago (Kya) during the Günz complex. Then, all lineages appear to have experienced a decline around the beginning of Upper Pleistocene (200 - 100 Kya), coinciding with a long glacial period (fig. 5C).

## Discussion

The population- and phylogenomic analyses of common wall lizards provide insights into how geological and climatic change in the Mediterranean Basin has shaped the evolutionary history of the Mediterranean fauna. We found strong support for a series of diversification events, range shifts, and extensive inter- and intraspecific gene flow taking place over the past six million years, connected to key events in the Mediterranean history (e.g. Cavazza and Wezel 2003; Hewitt 2011a).

Biogeographic and demographic analyses suggest that the MRCA of *P. muralis* stemmed from the Italian Peninsula. While molecular dating is fraught with difficulty, the timing of first lineage divergence corresponds well with the onset of the Messinian salinity crisis (Krijgsman et al. 1999, 2010; Duggen et al. 2003). This suggests that the divergence may have been triggered by declining sea levels and increased land connection, although aridification might have restricted suitable habitats (Jiménez-Moreno et al. 2010; Fiz-Palacios and Valcárcel 2013). The climatic changes that are associated with the end of the Messinian salinity crisis may have promoted allopatry and genetic differentiation that eventually resulted in the three distinct lineages on the Italian peninsula (CI, SI and SA).

The descendants of this ancient Italian assemblage appear to have followed an “Out-of-Italy” dispersal route to the Balkan and Iberian Peninsulas after the Messinian salinity crisis, a pattern that is consistent with dating of *Podarcis* fossil remains from Germany (Böttcher 2007). Both the RAD-Seq and WGS data supported six major genetic lineages for *P. muralis,* each with a distinct geographic distribution, which demonstrates the influence of the Iberian, Italian, and Balkan peninsulas on diversification within the Mediterranean Basin (e.g., Schmitt 2007; Hewitt 2011a). These lineages were largely discordant with the phylogenetic relationship based on mitochondrial genome data (Gassert et al. 2013; Salvi et al. 2013; Jablonski et al. 2019). Populations from the Iberian Peninsula, France and western Germany belong to the WE lineage (that can be further separated into north and south), and populations from the Balkan Peninsula can be assigned to two closely related lineages in the Southern (SB) and Northern (NB) Balkan. The contemporary distributions of the other three major lineages, Southern Alps (SA), Central Italy (CI), and Southern Italy (SI) are essentially restricted to Italy.

These lineages diverged between 6.24 and 2.5 Mya, but gene flow between lineages appears to have been extensive. This explains inconsistencies in estimates of relationships between *P. muralis* and other *Podarcis* species as well as between *P. muralis* lineages (Andrade et al. 2019; Salvi et al. 2021; Yang et al. 2021). These levels of ancient gene flow are consistent with studies of contemporary hybridization within *P. muralis*, which demonstrate that significant parts of a genome can introgress under positive selection (While et al. 2015; Yang et al. 2018; see also Schulte et al. 2012; Beninde et al. 2018). It is also consistent with estimates of ancient gene flow between extant lineages of *Podarcis* (Caeiro-Dias et al. 2020; Yang et al. 2021), which suggests that reproductive isolation evolves slowly in wall lizards. Nevertheless, the hybrid zones between different *P. muralis* lineages appear to be narrow and steep (e.g., Yang et al. 2020), and experimental studies suggest some degree of pre-mating reproductive isolation (Heathcote et al. 2016; MacGregor et al. 2017). Partial reproductive isolation is not unexpected given that some of the extant lineages have been separated for about five million years. The many hybrid zones, involving lineages of different age of divergence, makes *P. muralis* a useful system to study the evolution of pre- and post-copulatory mechanisms of speciation (e.g., Heathcote et al. 2016; Yang et al. 2020).

The most extensive introgression between *P. muralis* lineages involved the Southern Italy lineage, and explains the ambiguity with respect to its placement in the phylogenetic analysis. The SI lineage genome turned out to be a mosaic, comprising of roughly one third of alleles from the CI lineage in central Italy and two thirds of alleles from an early-diverged ancient lineage. We further revealed that this ancient lineage has contributed a substantial amount of genetic material to the MRCA of several extant lineages of *P. muralis*. These results demonstrate the value in complementing phylogenetic trees with introgression analyses of genomic data for detecting cryptic events in evolutionary histories (Than and Nakhleh 2009; Ottenburghs et al. 2016, 2017).

The WGS data also revealed that some lineages of *P. muralis* experienced ancient introgression from other *Podarcis* species. The WE lineage has received a substantial part of its genome from the MRCA of the Iberian species group (i.e., the *P. hispanicus* complex; see also Yang et al. 2021). There was also evidence for introgression from *P. siculus* into the SA and CI lineages on the Italian peninsula, although the signal was weak. More extensive genomic data is needed to establish if hybridization between *P. muralis* and local sympatric species has been more widespread and the extent to which is it ongoing.

Overall, the genetic structure revealed by genomic data is strongly discordant with the sub-species division of traditional taxonomy based on morphology (Gruschwitz and Böhme 1986; Biaggini et al. 2011) and molecular phylogenies based on mtDNA (Bellati et al. 2011; Gassert et al. 2013; Salvi et al. 2013; this study). Mito-nuclear discordance is common in nature (e.g. Zink and Barrowclough 2008; Toews and Brelsford 2012; Ivanov et al. 2018), and can result from several different processes, including incomplete lineage sorting (Firneno et al. 2020), introgression (Phuong et al. 2017; Ivanov et al. 2018), and sex-biased dispersal (Dai et al. 2013).

Some instances of mito-nuclear discordance in *P. muralis* are likely the result of introgression. This is particularly well illustrated by the NB, SA and WE lineages, where genomic data support exchange of mtDNA through introgression events between the three lineages, as well as with the MRCA of extant Iberian *Podarcis* species (Yang et al. 2021). Other instances of mito-nuclear discordance are better explained by incomplete lineage sorting. For example, the individual from San Remo, situated on the Italian coast close to the border to France, belonged to the SA lineage according to nuclear data but formed the sister clade to all *P. muralis* in the mitochondrial phylogeny. Since there was no signal of mtDNA or nuclear genomic introgression from closely related species, or evidence of introgression from a “ghost lineage”, this discordance appears to be a remnant of an ancient mtDNA that persisted throughout the evolutionary history of the SA lineage. Judging from the evidence for mito-nuclear discordance in other animals (e.g., Firneno et al. 2020), situations like these are probably not unusual but alternative hypotheses can be difficult to rule out (Toews & Brelsford 2012). Indeed, the genetic structure of *P. muralis* in this geographic region (south-western arc of the Alps) is poorly studied, and it is possible that more extensive sampling will reveal nuclear genomic signatures of an extinct lineage. Our data also supported a divergent mtDNA clade on the island of Elba (Bellati et al. 2011), but there was no evidence that this reflects a deep genetic divergence with mainland CI populations or an introgression event. Thus, incomplete lineage sorting of mtDNA haplotypes during or following isolation on Elba may best explain this discordance.

In the Mediterranean Basin, the Pliocene-Pleistocene climatic oscillations have been considered to play a key role for the current patterns of biodiversity and biogeography of animal species (reviewed in Hewitt 2000; Hewitt 2004). In particular, the genetic structure of many species has been explained by cycles of glacial contraction to refugia in southern peninsulas and inter-glacial expansion to northern regions (Hewitt 1996, 2000, 2004; Provan & Bennet 2008; Taberlet et al., 1998). Our results are consistent with this hypothesis, but suggest that all the major extant lineages were already present at the onset of the Quaternary climatic oscillations. Glacial cycles therefore appear to have played a less important role in initiating lineage divergence than might be assumed (reviewed in Hewitt 2004, 2011a), but nevertheless have been crucial for dictating the subsequent dynamics of these lineages. Indeed, our demographic simulations indicated that all lineages were affected by glacial cycles, including a population expansion (ca. 0.7 Mya) during the Günz complex and a severe decline (ca. 0.2 - 0.1 Mya) during the Riss glaciation. However, the inferred population dynamics differed somewhat between lineages. This suggests that lineages may have largely persisted in distinct regions during glacial and inter-glacial periods, including in northern refugia in France and eastern Europe, thereby promoting further genetic differentiation and reproductive isolation. The presence of several refugia within each of these regions might explain the high number of low-divergent mitochondrial sub-lineages found in Mediterranean and extra-Mediterranean regions (Salvi et al., 2013; Jablonski et al. 2019; see also fig. 1D). A similar pattern of mtDNA lineages has been found for other Mediterranean taxa, for example butterflies (Dincă et al. 2019; Hinojosa et al. 2019) and slow worms (Gvoždík et al. 2013). Allopatric isolation during glaciation may have also promoted reproductive isolation, and therefore contributed to the persistence of lineages following secondary contact, and explain the location and apparent stability of contemporary hybrid zones (e.g., between the SA and CI lineages; Yang et al., 2018, 2020).

In summary, the range-wide genomic approach employed in our study allowed a disentangling of the evolutionary history of a broadly distributed Mediterranean lizard species in unprecedented detail. We reveal an “Out-of-Italy” origin of the species about five Mya, roughly coinciding with the end of the Messinian salinity crisis. The species diversified into distinct lineages soon after its expansion from the Italian peninsula. The genomes of these lineages carries the signature of several major introgression events and of extinct lineages as well as the demographic imprints left by range expansions and contractions associated with glacial cycles.

## Materials and methods

### Samples and data collection

We collected samples from a total of 55 locations, covering the previously suggested genetic structure within *P. muralis* (fig. 1A; Gassert et al. 2013; Salvi et al. 2013). Detailed information for these samples is provided in supplementary table S1, and the collection permits are given in supplementary table S6. For each sample, we extracted total genomic DNA using the DNeasy blood and tissue kit (Qiagen, USA). We genotyped samples from all sites using RAD-Seq (deposited in NCBI Short Reads Archive [SRA] with accession number PRJNA486080). The RAD-Seq libraries were prepared following the protocol in Peterson et al. (2012) with modifications described in Yang et al. (2018). In addition, we conducted whole genome sequencing (WGS) for 16 individuals (supplementary table S2; supplementary fig. S1) with insert size of 300-500 bp on the Illumina HiSeq X platform by NOVOGENE Ltd. (Hong Kong). One individual of *P. bocagei*, *P. siculus*, and *P. tiliguerta* served as outgroups for all analyses and, in addition, *Archaeolacerta bedriagae* was included as outgroup for analyses involving WGS data. We also included WGS of 13 additional *Podarcis* species/lineages (accessible in NCBI under the accession number PRJNA715201; supplementary table S3), representing all species groups in *Podarcis* according to Yang et al. (2021), to investigate the gene flow between *P. muralis* and other *Podarcis* species.

### Data processing for sequencing data

We used STACKS (version 2.4; Catchen et al. 2011; Rochette et al. 2019) to process RAD-Seq reads and infer single nucleotide variants (SNVs) for each individual. At first, the “process_radtags” module was used to remove reads with low-quality scores (Phred score < 30), ambiguous base calls, or incomplete barcode or restriction site. Clean reads were mapped to the genome of *P. muralis* (Andrade et al. 2019) using BWA (Li and Durbin 2009). We used sorted bam files as input for the reference-based STACKS pipeline that contains modules “gstacks” and “populations” to estimate SNVs using a Marukilow model (Maruki and Lynch 2017). We also aligned WGS reads to the *P. muralis* genome using BWA (Li and Durbin 2009). We called SNVs and short indel variants using the GATK best practice workflow (DePristo et al. 2011). Only SNVs from autosomes were used in the following phylogeographic analyses.

### Mitochondrial genome

We assembled the mitochondrial genomes from WGS reads using NOVOPasty (Dierckxsens et al. 2017). The mitochondrial genome of *P. muralis* (accession FJ460597 from MitoZoa; D’Onorio et al. 2012) was set as a starting reference. A total of 6 Gb sequence reads from each sample were randomly extracted for the baiting and iterative mapping with default parameters. We aligned mitochondrial DNA (mtDNA) sequences using MUSCLE v3.8.31 (Edgar 2004). We excluded all ambiguous regions from the analyses to avoid false hypotheses of orthology.

### Population structure analysis

We inferred the genetic relationship between the samples based on genotypes from RAD-Seq data. A principal component analysis (PCA) was conducted in Plink (version 1.9; Chang et al. 2015) based on pairwise genetic distance. In addition, the population structure was inferred assuming different numbers of clusters (K) from 1 to 15 in ADMIXTURE (version 1.3.0; Alexander et al. 2009). We used 10-fold cross-validation (CV) to compare different numbers of clusters, in which the lowest CV value indicates the most likely number of clusters.

### Phylogenetic analysis

We reconstructed phylogenetic trees with maximum likelihood (ML) inference using IQ-TREE (Nguyen et al. 2015). We concatenated all SNVs generated from RAD-Seq dataset and inferred the phylogenies under a GTR+ASC model and 1,000 iterations of bootstrap replicates. We excluded 12 samples that showed admixed patterns in ADMIXTURE analysis (admixed ancestry > 10%). We also performed the same concatenation approach on SNVs for the 16 individuals with WGS data. An individual from KOP was excluded in the downstream analyses due to an admixed pattern. In addition, we used the “multispecies coalescent” approach implemented in ASTRAL-III (Zhang et al. 2018) to infer the phylogenetic relationships based on local trees of 200 kb fixed windows across the whole genome.

We inferred phylogenetic trees based on mitogenomic data implementing the highest-ranked model with 1,000 bootstrap replicates using IQ-TREE (Nguyen et al. 2015). We performed model selection for 1/2, and 3 codon positions for protein coding genes, and tRNA/rRNA genes with the setting “-m mf”.

### Gene flow analysis

We used D statistics (ABBA-BABA test; Patterson et al. 2012) to estimate the gene flow between both inter- and intraspecific lineages using “qpDstat” in AdmixTools (Patterson et al. 2012), using *A. bedriagae* as outgroup. We tested the significance level of D statistics through a block-jackknifing approach as implemented in AdmixTools, in which the z-score > 3.3 is considered significant (Patterson et al. 2012). First, we tested for interspecific introgression, between each of the major *P. muralis* lineages and geographically overlapping *Podarcis* species. We used the WGS data for 13 *Podarcis* species and lineages as “target_taxon” in the test (*muralis*_A, *muralis*_B, target_taxon, outgroup). Second, we also estimated the intraspecific gene flow between the major *P. muralis* lineages (i.e., muralis_A, muralis_B, muralis_C, outgroup).

We further conducted phylogenetic network analysis using phyloNet (Wen et al. 2008) to infer reticulation events between the *P. muralis* lineages (i.e., intraspecific introgression). *P. bocagei* was also included in this analysis to represent the Iberian *Podarcis* species, since we identified a strong signal of introgression in interspecific D statistics and in a previous study (Yang et al. 2021). We made use of high-quality local trees derived from 200 kb windows with mean bootstrap > 80, and extracted 1,000 random trees per run with 10^6^ chain-length and 50% burn-in length in MCMC_gt module. We performed 100 independent runs, extracted all output networks with more than 50% posterior probability, and summarized the results by generating a correlation matrix of those networks based on Luay Nakhleh’s metric of reduced phylogenetic network similarity (Edelman et al. 2019).

Based on the phylogenetic network, we used the program “qpGraph” from ADMIXTOOLS (Patterson et al. 2012) to fit the evolutionary history for all *P. muralis* lineages while accounting for introgression. qpGraph optimizes the fit of a proposed admixture graph in which each node can be descended either from a mixture of two other nodes, or from a single ancestral node. We calculated the proportion of introgressed alleles ted by f4-ratio tests (Patterson et al. 2012). To identify the genomic regions with signatures of introgression, we calculated the absolute genetic divergence (Dxy) between lineage pairs, and the fd statistics based on 200 kb windows across the genome. A significantly low Dxy and high fd identify an introgressed genomic region tested by 1,000 permutations.

### Divergence time estimation

We performed divergence time analysis between *P. muralis* lineages based on the WGS phylogeny using the MCMCtree program in the PAML package (Yang 2007). According to the fossil-calibrated Lacertini phylogeny of Garcia-Porta et al. (2019), the divergence time between *Podarcis* and other closely related clades was estimated at 37.55 million years ago (Mya), and the crown node of *Podarcis* species was at 18.60 Mya. We specified these calibration constraints with soft boundaries by using 0.025 tail probabilities above and below the limit in the built-in function of MCMCtree. To exclude the confounding effect of introgression events on topology and divergence time estimates, we only retained those genomic regions (346 windows with length of 200 kb) whose local trees were consistent with the consensus phylogeny. The independent rate model (clock = 2) was used to specify the rate priors for internal nodes. The MCMC run was first executed for 10^7^ generations as burn-in and then sampled every 150 generations until a total of 100,000 samples were collected. We compared two MCMC runs using random seeds for convergence, which yielded similar results.

### Biogeographic and demographic analysis

We used BioGeoBEARS (Matzke 2013) to infer the possible ancestral range of *P. muralis* and the number and type of biogeographical events dispersal leading to the distribution of extant lineages. We defined seven biogeographic areas covering the current distribution of this species: Iberian Peninsula, Western Europe, Southern Alps, Central Italy, Southern Italy, Northern Balkans and Southern Balkans. The time-calibrated phylogeny for *P. muralis* lineages was applied under three models - dispersal extinction cladogenesis (DEC), dispersal vicariance analysis (DIVALIKE), and Bayesian inference of historical biogeography for discrete areas (BAYAREALIKE). We selected the best-fitting model for comparisons among models based on AICc.

To reconstruct the detailed demographic history of each lineage, we applied the pairwise sequential Markovian coalescence (PSMC) model (Li and Durbin 2011) with the following parameters “-N25 -t15 -r5 -p4+25*2+4+6”. We selected two to three sequenced individuals from each extant lineage. We excluded the SI lineage in this analysis due to its strikingly heterogeneous genome. On the basis of the time-calibrated phylogeny, we made use of a mutation rate of 1.98 × 10^−9^ mutations per site per year. The generation time for *P. muralis* was set to 2 years (Barbault and Mou 1988).

## Data Availability

All sequence data generated in this study have been deposited in NCBI Sequence Reads Archive (SRA) with accession number PRJNA486080 (RAD-Seq data) and PRJNA715201 (whole genome data).

## Acknowledgements

Computations were performed on resources provided by SNIC through the center for scientific and technical computing at Lund University under Project SNIC 2017/4-39, and Uppsala Multidisciplinary Center for Advanced Computational Science under Project SNIC 2017/5-8 and Uppstore 2018/2-18. This study was funded by the Swedish Research Council (E0446501), the Crafoord Foundation (20160911), the National Geographic Society, the British Ecological Society, the Royal Society of London, and a Wallenberg Academy Fellowship to TU, starting grants from the Swedish Research Council (2020-03650) and the European Research Council (948126) to NF, a LabEx TULIP grant (ANR-10-LABX-41) to FA and the Slovak Research and Development Agency (APVV-19-0076) to DJab.

