## Supplementary materials for "Population genomics of wall lizards reflects the dynamic history of the Mediterranean Basin"

Supplementary materials contain:

6 supplementary tables

8 supplementary figures

**Supplementary table S1** Information of sampling sites for RAD-Seq samples in this study.

| Site | Longitude | Latitude |
|---|---|---|
| ALB2 | 19.633 | 42.3685 |
| ALB4 | 19.668 | 40.9833 |
| AQF | 16.242 | 39.9874 |
| BUL_MT | 27.475 | 41.9706 |
| BUL4 | 24.81 | 42.64 |
| BUL5 | 25.99 | 41.76 |
| BV | 11.539 | 45.4106 |
| CAR | -3.095 | 47.5746 |
| CCR | 14.404 | 44.9586 |
| CDG | 14.071 | 41.9954 |
| CE | 11.291 | 44.7315 |
| COR | 12.228 | 42.7033 |
| CR | 10.565 | 43.5748 |
| CRO | 17.053 | 43.3423 |
| CRS | 14.955 | 40.7835 |
| CV | 11.936 | 43.6981 |
| CX | 10.341 | 44.5331 |
| EF_W | 6.89 | 49.99 |
| ENF | 10.263 | 42.8277 |
| FRM | 13.741 | 43.172 |
| GRU | 9.1654 | 45.5682 |
| KOP | 13.727 | 45.5488 |
| LE | 9.6107 | 44.1686 |
| MG | 10.162 | 44.0122 |
| MON | -1.8178 | 41.7902 |
| MOR | -3.8295 | 40.8274 |
| MP | 11.163 | 44.0926 |
| NL | 8.4142 | 44.2063 |
| NM | 2.6762 | 48.2879 |
| OLY | 22.257 | 40.1155 |
| OV | 12.934 | 42.13 |
| PDT | -3.8351 | 47.7966 |
| PEN | -0.3339 | 40.2481 |
| PI | 9.5029 | 44.8198 |
| PY | 1.1052 | 42.8595 |
| PZ | -0.8392 | 46.7844 |
| RCM | 14.35 | 41.4924 |
| RO | 12.544 | 41.8586 |

| | | |
|---|---|---|
| ROM2 | 21.68 | 44.73 |
| ROM7 | 21.72 | 46.11 |
| ROM9 | 23.96 | 45.42 |
| SER_BO | 20.655 | 43.4295 |
| SER1 | 20.45 | 44.79 |
| SER3 | 19.77 | 43.46 |
| SER4 | 19.58 | 43.85 |
| SER7 | 22.52 | 42.43 |
| SLOVK2 | 17.073 | 48.1441 |
| SLOVK4 | 19.483 | 49.1017 |
| SR | 7.7729 | 43.8183 |
| ST | 9.9046 | 44.0823 |
| TRC | 21.466 | 38.5896 |
| TYG | 22.362 | 36.9519 |
| UR | 12.636 | 43.7236 |
| VI | 10.263 | 43.8427 |
| VTS | 21.388 | 40.6138 |

**Supplementary table S2** Information of sampling sites for WGS in this study

| Site | Longitude | Latitude |
|---|---|---|
| AQF | 16.24217 | 39.98745 |
| Elba_east | 10.360371 | 42.808416 |
| Elba_west | 10.169758 | 42.786448 |
| GRC | 21.726 | 39.1825 |
| KOP | 13.724 | 42.5488 |
| LE | 9.610654 | 44.16861 |
| NL | 8.414208 | 44.20628 |
| NM | 2.67624 | 48.28794 |
| PDT | -3.83513 | 47.79656 |
| PEN | -0.67 | 40.37 |
| RCM | 14.34966 | 41.49243 |
| RO | 12.54419 | 41.85863 |
| SLOVK | 17.075601 | 48.143695 |
| SR | 7.772901 | 43.81833 |
| TK | 27.42 | 41.97 |
| TU | 10.720143 | 43.544483 |

**Supplementary table S3** *Podarcis* species and outgroup used in gene flow analysis

| Sample_ID | Species | Note |
|---|---|---|
| Ab387.8 | *Archaeolacerta bedriagae* | Outgroup |
| PB01 | *Podarcis bocagei* | |
| DB28361 | *Podarcis erhardii* | |
| 2100-110 | *Podarcis filfolensis* | |
| DB7019 | *Podarcis hispanicus* | PHGAL lineage |
| DB28386 | *Podarcis levendis* | |
| DB20801 | *Podarcis melisellensis* | |
| DB13118 | *Podarcis pityusensis* | |
| CAP3 | *Podarcis siculus* | North lineage (N) |
| TRC3 | *Podarcis siculus* | South lineage (S) |
| DB16305 | *Podarcis tauricus* | |
| 395A | *Podarcis tiliguerta* | Corsica lineage (C) |
| PF2 | *Podarcis tiliguerta* | Sardinia lineage (S) |
| DB26449 | *Podarcis waglerianus* | |

**Supplementary table S4** Results of D statistics for intraspecific comparison for *P. muralis* lineages

| W | X | Y | D | Zscore |
|---|---|---|---|---|
| Southern Balkan | Northern Balkan | Central Italy | -0.048 | -13.511 |
| Western Europe (N) | Northern Balkan | Central Italy | -0.143 | -31.426 |
| Western Europe (N) | Southern Balkan | Central Italy | -0.1087 | -22.867 |
| Western Europe (S) | Northern Balkan | Central Italy | -0.1215 | -24.86 |
| Western Europe (S) | Southern Balkan | Central Italy | -0.086 | -16.76 |
| Western Europe (S) | Western Europe (N) | Central Italy | 0.0365 | 10.546 |
| Western Europe (S) | Western Europe (N) | Northern Balkan | 0.0381 | 9.514 |
| Central Italy | Northern Balkan | Southern Alps | -0.0783 | -29.906 |
| Central Italy | Southern Italy | Southern Alps | 0.1927 | 94.085 |
| Central Italy | Southern Balkan | Southern Alps | -0.0387 | -13.885 |
| Southern Italy | Northern Balkan | Southern Alps | -0.2422 | -100 |
| Southern Italy | Southern Balkan | Southern Alps | -0.2122 | -79.013 |
| Southern Balkan | Northern Balkan | Southern Alps | -0.0479 | -16.372 |
| Western Europe (N) | Central Italy | Southern Alps | -0.0274 | -6.141 |
| Western Europe (N) | Northern Balkan | Southern Alps | -0.1021 | -26.528 |
| Western Europe (N) | Southern Italy | Southern Alps | 0.1463 | 36.376 |
| Western Europe (N) | Southern Balkan | Southern Alps | -0.0668 | -16.827 |
| Western Europe (S) | Central Italy | Southern Alps | -0.0147 | -3.089 |
| Western Europe (S) | Northern Balkan | Southern Alps | -0.0907 | -21.724 |
| Western Europe (S) | Southern Italy | Southern Alps | 0.1601 | 35.693 |
| Western Europe (S) | Southern Balkan | Southern Alps | -0.0543 | -12.955 |
| Western Europe (S) | Western Europe (N) | Southern Alps | 0.0205 | 5.708 |
| Central Italy | Northern Balkan | Southern Italy | 0.1074 | 52.96 |
| Central Italy | Southern Balkan | Southern Italy | 0.1071 | 41.563 |
| Southern Balkan | Northern Balkan | Southern Italy | -0.0033 | -1.27 |
| Western Europe (N) | Central Italy | Southern Italy | -0.1806 | -53.737 |
| Western Europe (N) | Northern Balkan | Southern Italy | -0.09 | -23.136 |
| Western Europe (N) | Southern Balkan | Southern Italy | -0.089 | -23.198 |
| Western Europe (S) | Central Italy | Southern Italy | -0.1666 | -47.366 |
| Western Europe (S) | Northern Balkan | Southern Italy | -0.0727 | -17.622 |
| Western Europe (S) | Southern Balkan | Southern Italy | -0.0717 | -17.425 |
| Western Europe (S) | Western Europe (N) | Southern Italy | 0.0276 | 8.129 |
| Western Europe (S) | Western Europe (N) | Southern Balkan | 0.0399 | 9.492 |
| Southern Balkan | Northern Balkan | Western Europe (N) | 0.0298 | 7.692 |
| Southern Balkan | Northern Balkan | Western Europe (S) | 0.0316 | 7.471 |

Topology for D statistics is (W, X, Y, O), in which the O is the constant ougroup *A. bedriagae*.

**Supplementary table S5** Model and parameters of ancestral range estimation of extant *Podarcis muralis* lineages

| Model | lnL | nParam | d | e | AICc | w |
|---|---|---|---|---|---|---|
| DEC | -27.55 | 2 | 1.29 | 3.77 | 60.09 | $4.5 \times 10^{-5}$ |
| DIVALIKE | -22.84 | 2 | 1.21 | $10^{-12}$ | 50.69 | 0.0049 |
| BAYAREALIKE | -41.02 | 2 | 1.64 | 5 | 87.03 | $6.3 \times 10^{-11}$ |

lnL: log-likelihood; d: rate of range expansion; e: rate of range reduction through extirpation in an area; AICc: Akaike information Criterion adjusted for small samples; w: Akaike weights.

**Supplementary table S6** Permits for specimen collection

| Permit ID | Issuing authority |
|---|---|
| No. 6584 | Ministry of Environment of Albania |
| 141911/1567/26-5-2016 and 73ΟΣ4653Π8-7Ψ8 | Ministry of Environment and Energy |
| ref2017-01666 and #2012-10 | Direction Départementale de la Protection des Populations of France |
| DPN-2009-0005106, PNM-2012-0009747 and PNM-2017-0008287 | Ministry of Environment of Italy |
| No. 9303/2009-2.1/jam and 4145/2011-2.2 | Ministry of the Environment of the Slovak Republic |
| 35601-66/2013-4 | Ministry of Agriculture and the Environment of Slovenia |
| OAEN/SVSIA/avp_10_169_aut | Organismo Autónomo Espacios Naturales de Castilla – La Mancha, Spain |
| EP/SA/373/2016 | Delegación Territorial de Salamanca, Junta de Castilla y León, Spain |
| 500201/24/2014/3218 | Departamento de Agricultura, Ganadería y Medio Ambiente, Gobierno de Aragón, Spain |
| B.30.2.DEU.O.A8.00.00/54 | Turkish Environmental Authority |
| Presidential Decree 67/81* | President of Greece |

* issued to the Natural History Museum of Crete

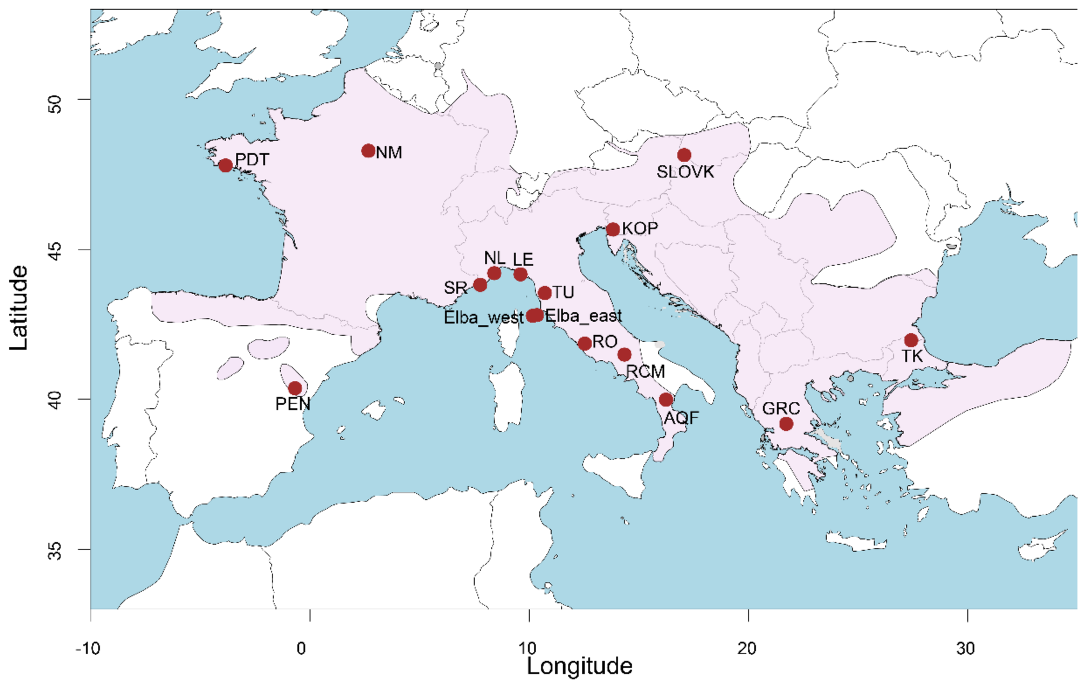


**Supplementary fig. S1** Collection sites for samples used for whole genome sequencing in this study. The pink area represents the distribution range of *P. muralis*.

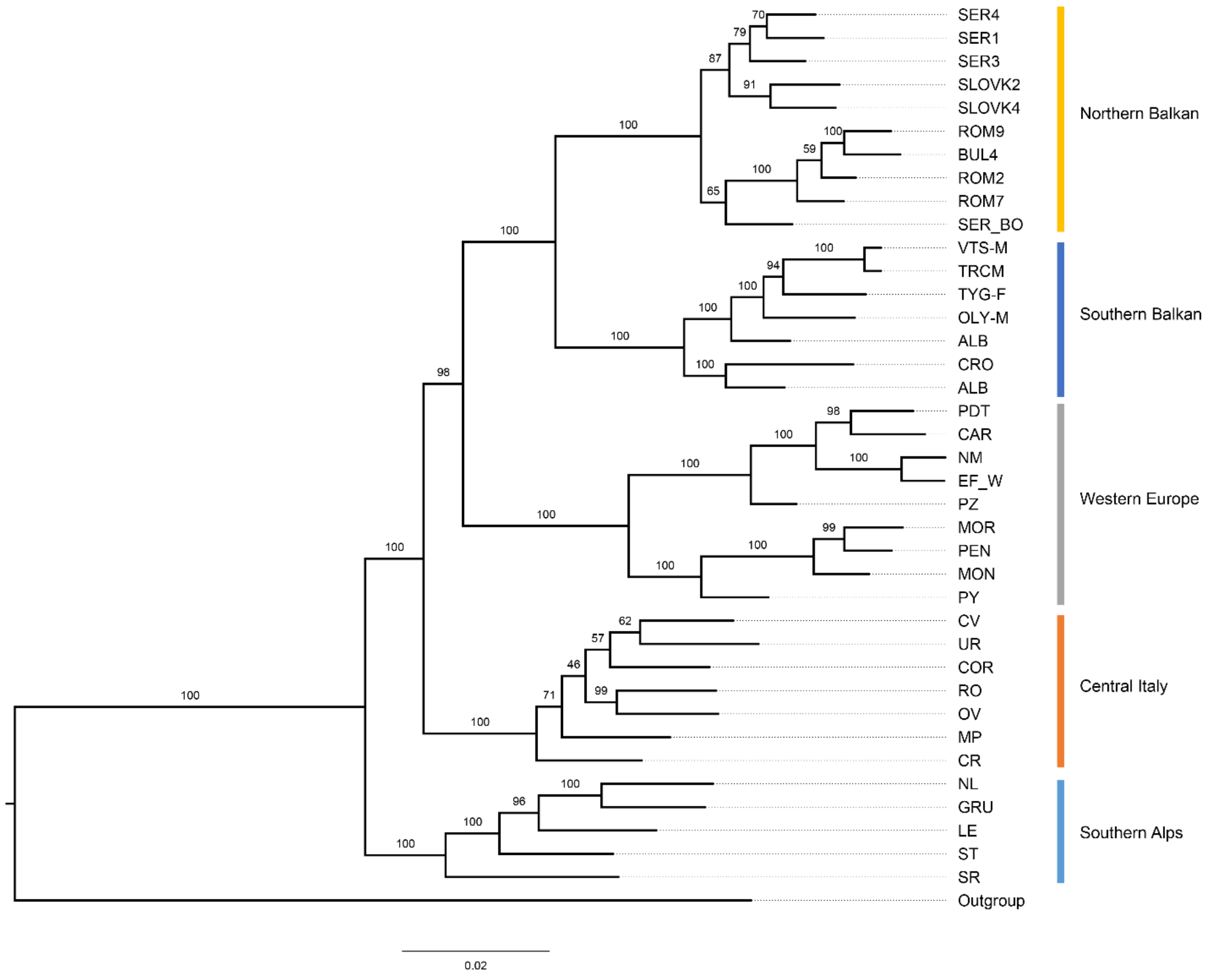


**Supplementary fig. S2** Maximum likelihood phylogeny inferred by RAD-Seq data without the Southern Italy lineage. The numbers above branches indicate the bootstrap support values.

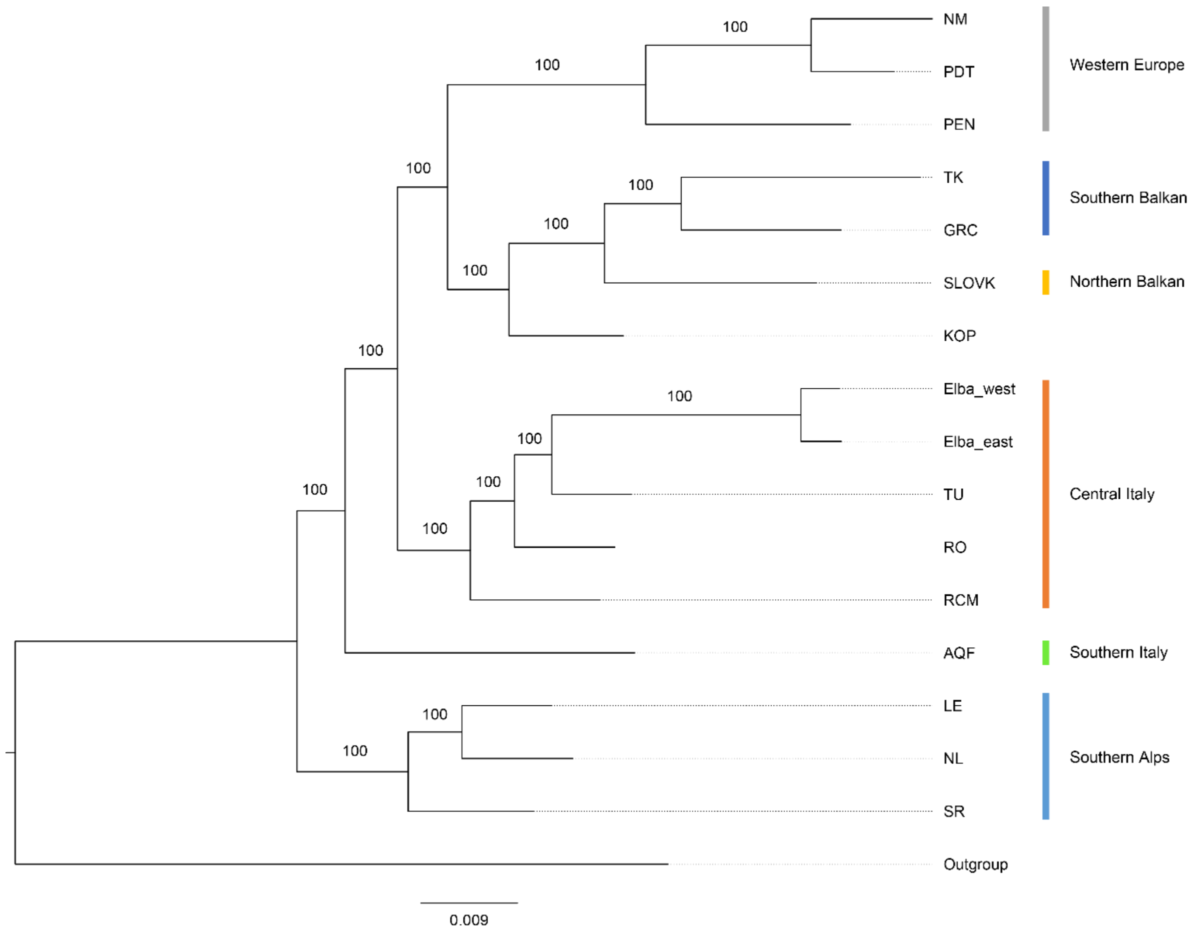


**Supplementary fig. S3** Maximum likelihood phylogeny inferred by WGS data for all 16 samples. The numbers above branches indicate the bootstrap support values. The admixed individual from KOP was sister to the Balkan clades.

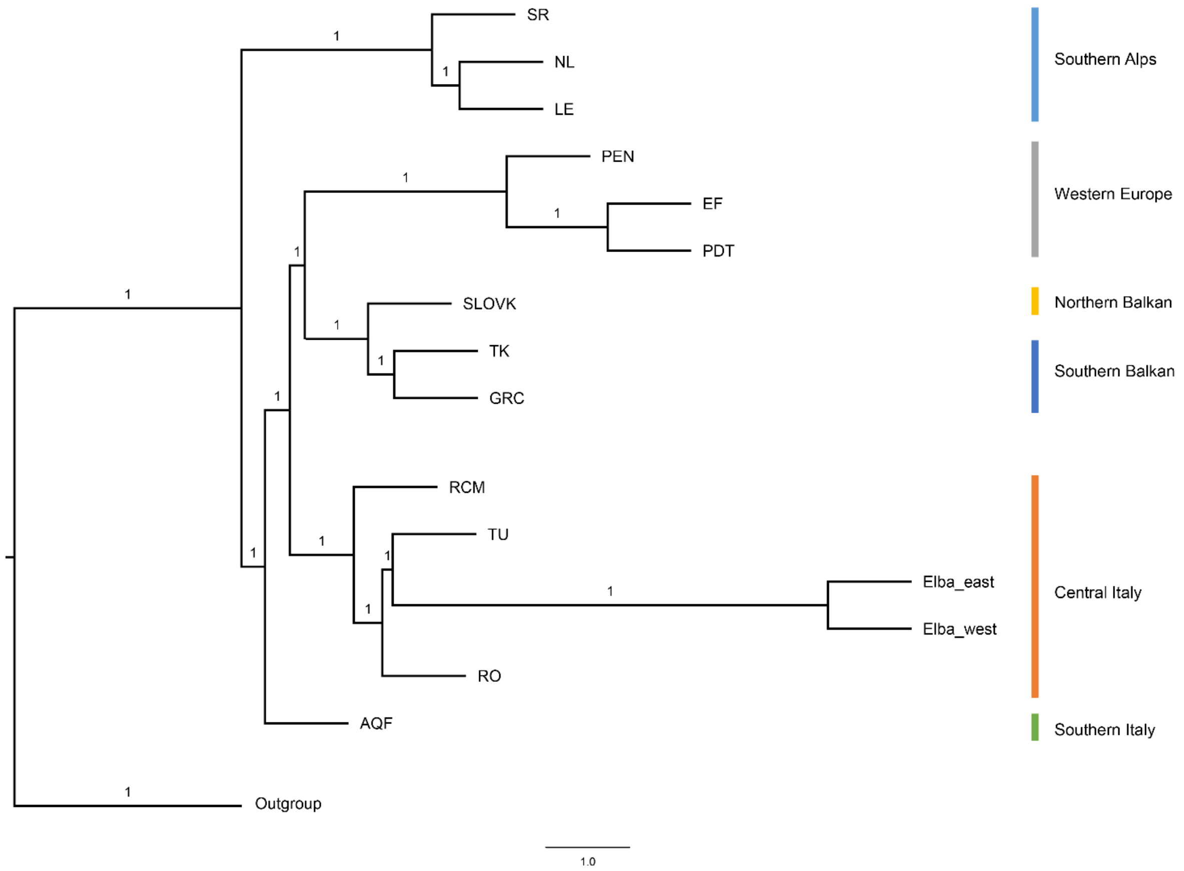


**Supplementary fig. S4** Phylogeny inferred by multispecies coalescent approach for whole genome sequencing data. The numbers above branches indicate the bootstrap values for genetic lineages.

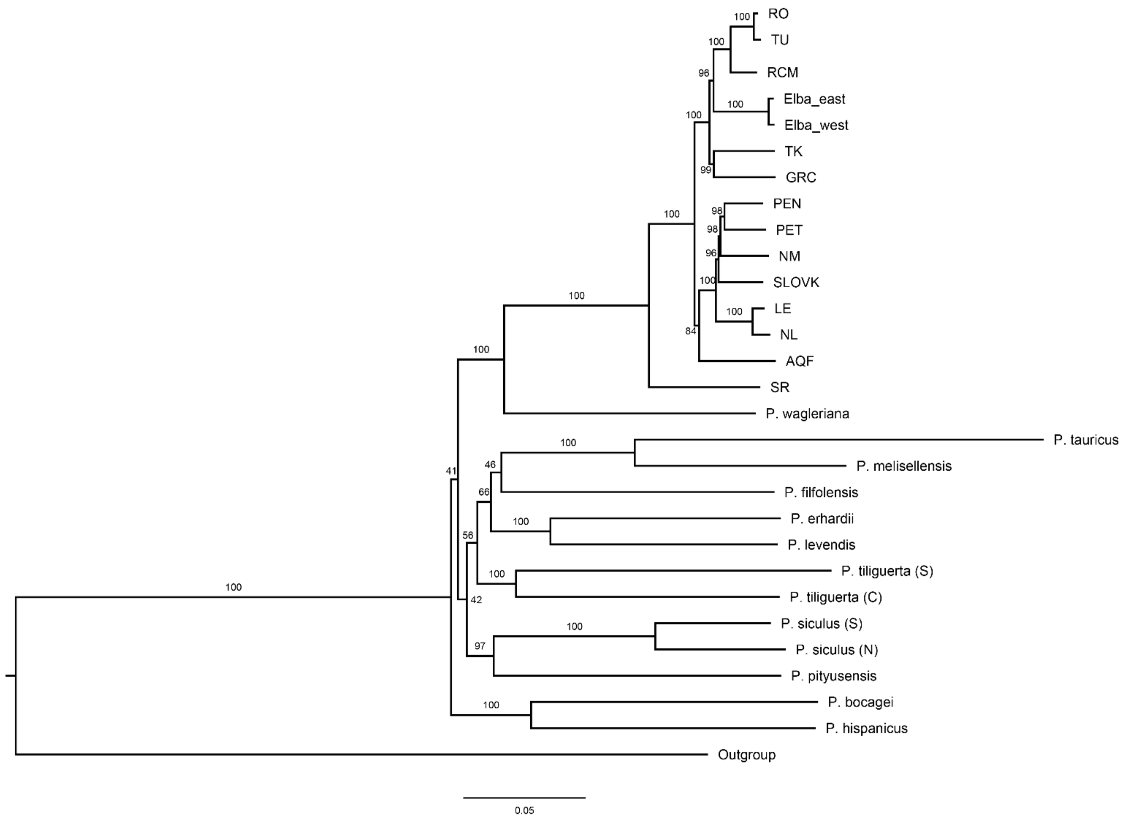


**Supplementary fig. S5** The phylogeny inferred by mitochondrial genome data with *Podarcis* samples. The numbers above branches indicate the bootstrap values for genetic lineages.

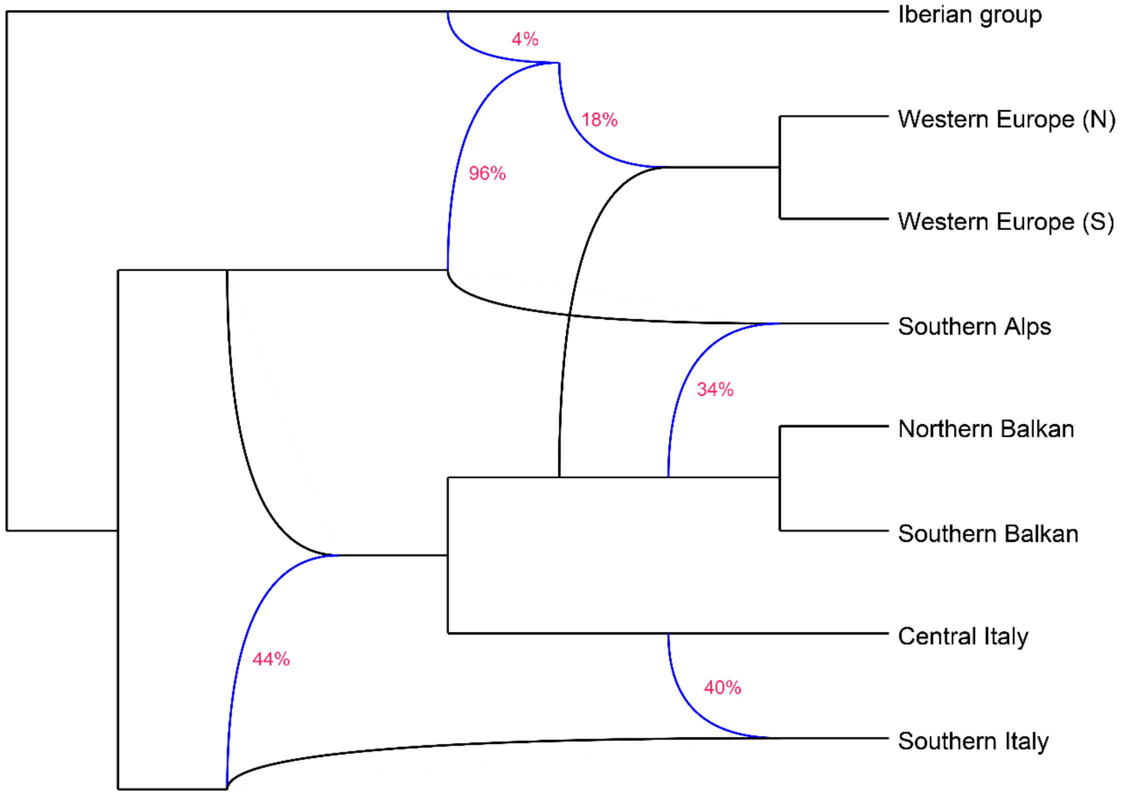


**Supplementary fig. S6** Phylogenetic network inferred by phyloNet. The blue lines represent the reticulation events, and the red numbers represent the proportion of alleles from parental lineages.

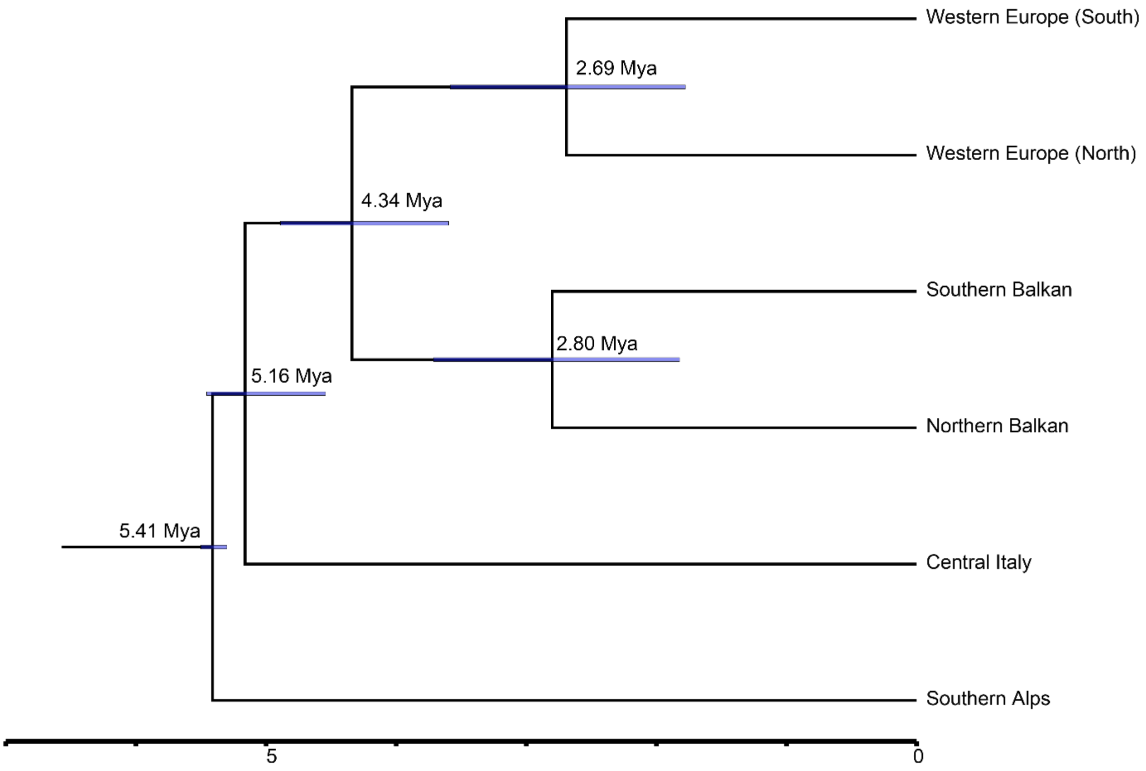


**Supplementary fig. S7** Time-calibrated phylogeny for *P. muralis* without the Southern Italy lineage. The numbers indicate the estimated ages for each node. The blue bars represent the confidence interval of nodes.

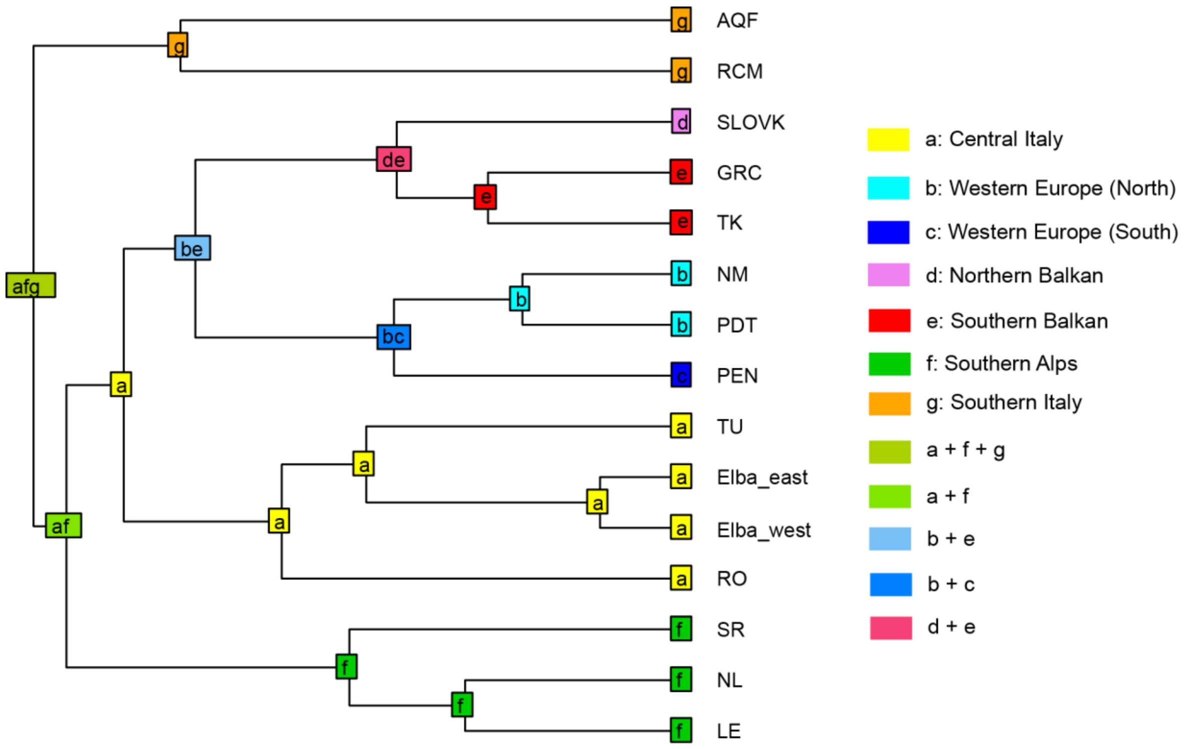


**Supplementary fig. S8** The best-fitting model (DEC) in BioGeoBEARS analysis for *P. muralis* lineages based on WGS data. The colored squares represent the distributed area.
